# Recursive Repeat Extender (RRE): A recursive approach to automatically extend repeat element models

**DOI:** 10.64898/2026.04.14.718546

**Authors:** Francisco Falcon, Elly M. Tanaka, Diego Rodriguez-Terrones

**Author notes:** These authors contributed equally to this work (Francisco Falcon, Diego Rodriguez-Terrones).

## Abstract

Repetitive elements, including transposable elements (TEs), are integral structural components of eukaryotic genomes; consequently, their identification and classification are crucial to their study. Several approaches have been developed to perform *de novo* genome-wide repeat identification through pairwise sequence comparisons; however, they often generate truncated repeat models due to their sampling strategies and the substantial fragmentation of many of the older repeat copies in the genome. To improve repeat models generated *de novo*, several algorithms have been developed that increase model length via the BEEA (BLAST-Extend-Extract-Align) approach, in which genomic instances of each repeat are identified with BLAST, their coordinates are extended, and a refined model is generated by aligning the extended sequences. Nevertheless, these extension algorithms exhibit two key limitations that hinder the reconstruction of highly degenerate and fragmented repeats: the use of BLAST as a search algorithm – which limits their sensitivity in detecting highly diverged sequences – and the use of a single search step, which precludes the reconstruction of extensively fragmented repeat models. In this work, we present a novel approach to extend repeat models, called RRE (Recursive Repeat Extender), which uses profile hidden Markov models (HMMs) to search for repeat elements with high sensitivity and employs a recursive extension strategy that iteratively searches and extends the repeat model, using the extended model from each round as input for the next and continuing until no additional sequence can be incorporated. We apply RRE to repeat libraries generated *de novo* from five model organisms, and our results show that RRE-generated repeat libraries contain fewer but longer repeat models and can identify a larger proportion of the genomes as repetitive than RepeatModeler2-generated repeat libraries. Notably, RRE can reconstruct highly degenerate repeats such as CR1_Mam, producing a model that achieves similar coverage to the reference Dfam model while extending it by an additional 131 bp that were not captured in the reference model. Overall, RRE enables the automatic improvement of *de novo* repeat libraries and the reconstruction of highly degenerate and fragmented repeats.

## Introduction

Repetitive elements are major structural components of eukaryotic genomes and based on their distribution within the genome they can be classified into tandem repeats and interspersed repeats. Among the interspersed repeats, transposable elements (TEs) are the most abundant type (Osmanski et al., 2023) and their expansion is a major determinant of genome size (Kapusta et al., 2017) and can extensively influence gene regulatory networks (Senft & Macfarlan, 2021). Because of their role in genome evolution, the identification and classification of transposable elements is essential in modern genomics.

Algorithms such as RepeatModeler2 (Flynn et al., 2020) and REPET (Flutre et al., 2011) identify repeats by first sampling genomic sequences and then comparing them using pairwise alignment tools such as BLAST. These alignments are then used to construct a graph in which nodes represent individual repeat instances and edges represent alignments between them (Bao & Eddy, 2002). After pruning the graph, these algorithms generate a repeat model that summarizes the sequence diversity of a repeat family (a group of repeats that share a common ancestor) by generating a multiple sequence alignment (MSA) from the genomic instances. The resulting collection of repeat models is commonly referred to as a repeat library. However, these approaches often generate incomplete or fragmented repeat models (i.e. a single repeat family may be represented by multiple partial models) (Baril et al., 2024; Flynn et al., 2020), largely due to a combination of their sampling strategies (i.e. incrementally sampling the genome until a certain amount has been sampled (Flynn et al., 2020)) and the progressive fragmentation of repeat instances over evolutionary time, driven in part by the insertion of younger transposable elements into pre-existing copies.

To refine fragmented repeat libraries generated by *de novo* approaches, algorithms such as EarlGrey (Baril et al., 2024), MCHelper (Orozco-Arias et al., 2024), and TEtrimmer (Qian et al., 2025) use a BLAST-Extract-Extend-Align (BEEA) approach to extend each repeat model. In short, these algorithms refine the repeat model by: i) searching each repeat model in a genome using BLAST (Basic Local Alignment Search Tool); ii) extending the resulting genomic coordinates by a given number of nucleotides (e.g. 250bp); iii) aligning the genomic sequences corresponding to these genomic coordinates in a multiple sequence alignment (MSA), which is used to generate a new repeat model. The process of extending genomic coordinates and aligning genomic sequences is repeated until the repeat model reaches a plateau in size. Extension algorithms increase the length and decrease the number of repeat models in a repeat library (Baril et al., 2024; Orozco-Arias et al., 2024; Qian et al., 2025). Nonetheless, these approaches are unable to reconstruct highly fragmented or degenerated repeats.

Repeat elements found in vertebrate genomes are ancient, representing remnants of transposable elements that were active millions of years ago (Smit, 1999). Because most repeat insertions evolve under neutral selection, they have accumulated numerous insertions, deletions, and point mutations over time (Arkhipova, 2018). Interestingly, some repeats exhibit particularly high levels of sequence divergence and fragmentation and are present across multiple clades, indicating that their transposition activity occurred hundreds of millions of years ago, preceding major speciation events. Their persistence over such long evolutionary timescales suggests that at least some of these elements have been co-opted by their hosts and retained due to functional roles. Despite considerable interest in the regulatory roles of TE-derived sequences, most research has focused on relatively young TEs and virtually nothing is known about these ancient TEs. In fact, out of the 1401 TE families identified to date in the human genome and listed on the Dfam repeat database, only 68 are annotated as being shared across mammals, 100 among amniotes and only 19 across tetrapods (Wheeler et al., 2012). This underscores the exceptional difficulty in identifying TEs dating back to these evolutionary intervals and highlights the need for specialized tools able to reconstruct them.

Given their likely importance in gene regulation, in this work we’ve set out to develop a computational approach to reconstruct these highly degenerate elements, which throughout this work we refer to as ‘ancient TEs’. While finding seeds that partially cover the most conserved regions of these ancient repeat families is possible with *de novo* methods such as RepeatModeler2, extension approaches that rely on the BEEA paradigm would fail to reconstruct a complete repeat model. The first impediment to reconstructing ancient repeats is the use of BLAST as a search engine, which relies on fixed-length exact k-mer matches (which can be as low as 7 nucleotides) and has a detection threshold near 70% identity when using its most permissive parameters (Altschul et al., 1990; States et al., 1991). As we show later in this work, this sensitivity is insufficient to detect the most degenerate regions of ancient TEs. The second hurdle for BEEA algorithms is the reliance on a single search step during the extension process. Because repeat elements accumulate insertions and deletions over time, loci rarely contain complete copies. Therefore, in ancient repeats, the search-once but extend multiple times approach of BEEA algorithms is only able to reconstruct the region adjacent to the seed model. To reconstruct these highly fragmented repeats, a recursive extension approach involving cycling between search and extension steps is required in order to progressively slide the reconstruction process along the length of the entire repeat and make use of all available genomic information.

In this work, we introduce Recursive Repeat Extender (RRE), which employs a novel recursive extension approach and profile HMMs (hidden Markov models) to extend repeat sequences in a genome, thereby overcoming the challenges of extending highly degenerate and fragmented repeats. To test RRE, we compared it against a *de novo* repeat annotation approach (RepeatModeler2) and are-implementation of the BEEA algorithm that uses HMMER instead of BLAST (HEEA: HMMER, Extract, Extend, and Align) in five species (*Caenorhabditis elegans, Drosophila melanogaster, Danio rerio, Mus musculus*, and *Homo sapiens*) whose repeat annotations have been extensively curated. Our results indicate that RRE’s recursive extension strategy outperforms the iterative extension strategy used in BEEA approaches, producing repeat libraries with fewer but longer repeat models and annotating a larger proportion of the genome as repetitive. Finally, to demonstrate that this approach can be used to extend ancient repeats, we use RRE to reconstruct a truncated version of the ancient mammalian repeat CR1_Mam, adding 131 bp missing from the reference model.

## Results

### HMMER outperforms BLAST in finding highly degenerate repeats

Conventional BEEA approaches rely on BLAST to identify repeat elements in a genome, but they are constrained by BLAST’s sensitivity limitations at high divergences, which stem primarily from its seed-based search strategy and alignment scoring scheme. An alternative to BLAST is HMMER, which uses profile hidden Markov models (HMMs). HMMs model the probability of each nucleotide occurring at each position, as well as gap and insertion probabilities, making them more sensitive at identifying highly degenerate sequences (Eddy, 2023). Although it has already been shown that HMMER improves overall repeat identification at the library level (Wheeler et al., 2012), its performance relative to BLAST for the detection of highly degenerate repeats has not been specifically evaluated.

To compare the performance of both methods, we analyzed the genomes of five species spanning a diversity of repeat landscapes: *Caenorhabditis elegans, Drosophila melanogaster, Danio rerio, Mus musculus*, and *Homo sapiens*. We employed repeat models from the Dfam database, which provides both BLAST-searchable consensus sequences and HMMER-searchable HMMs. Overall, HMMER outperformed BLAST when searching for repeat models in most genomes, identifying 37.4% more base pairs as repeats in *D. melanogaster*, 3.6% in *D. rerio*, 30.2% in *M. musculus*, and 2.3% in *H. sapiens*, but saw a decrease of 35.4% in *C. elegans*. At the individual repeat model level, HMMER outperformed BLAST in 79.1% of the models in *C. elegans*, 89.5% in *D. melanogaster*, 88.4% in *D. rerio*, 96.1% in *M. musculus*, and 90.7% in *H. sapiens* (Fig. 1A). Overall, these results are in agreement with previous observations indicating that HMMER is substantially more sensitive than BLAST when searching repeat models (Wheeler et al., 2012).

**Figure 1.**
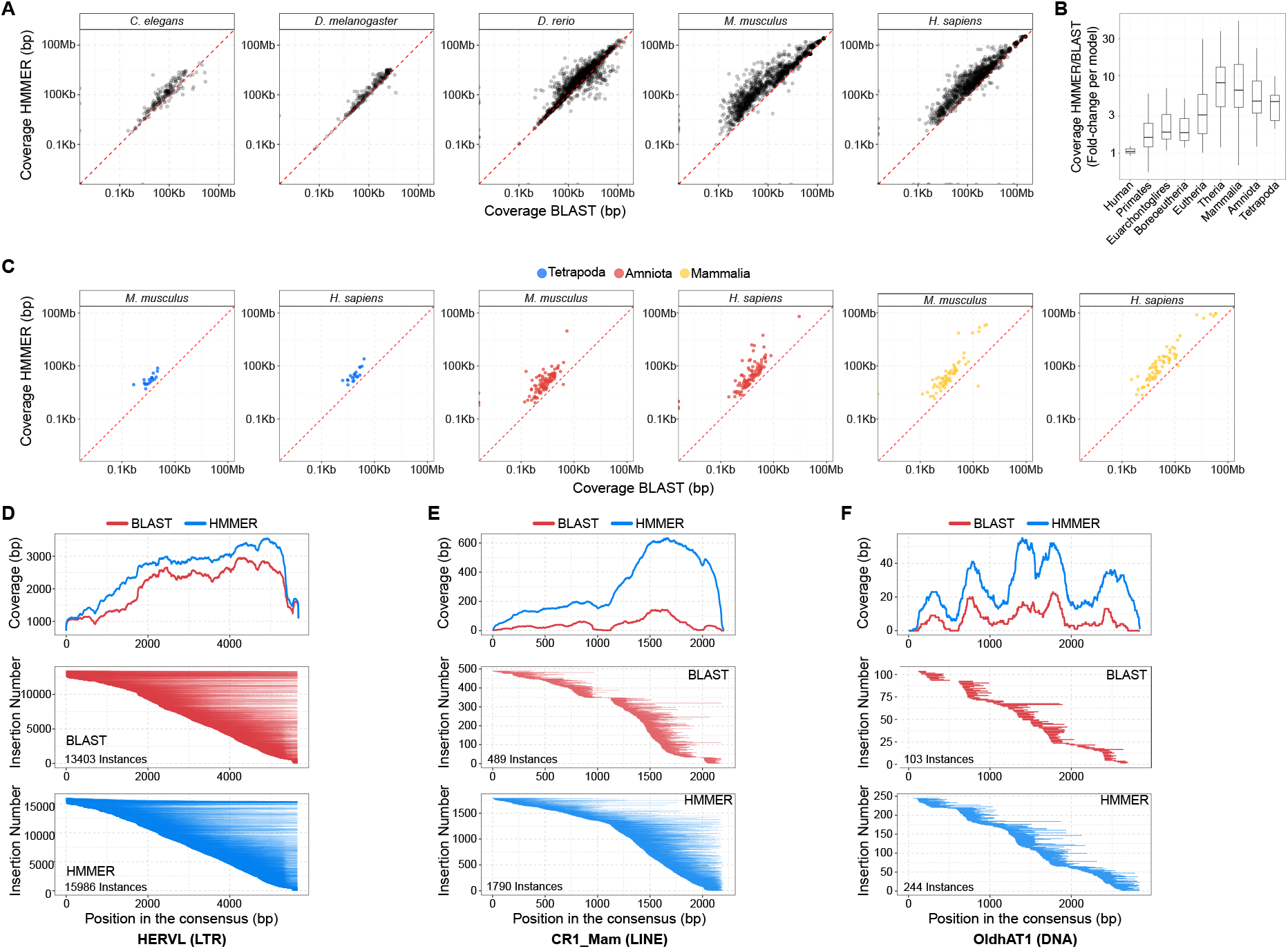
HMMER outperforms BLAST in identifying repeats. A) Repeat coverage (in bp) found with HMMER (filtered using the GA threshold provided in the Dfam models) and BLAST (filtering using e-value <=1e-05) for repeats in the Dfam database, where the red dotted line represents the identity line. B) Fold-change of the coverage between HMMER and BLAST per repeat model, grouped by the estimated transposition time listed in Dfam. C) Repeat coverage (in bp), for older repeats from the Tetrapoda, Amniota, and Mammalia lineages found in the Dfam database using HMMER (filtered using the GA threshold provided in the Dfam models) and BLAST (filtering using e-value <=1e-05). D-F) Coverage along the length of the repeat model and representation of the segments spanned by individual instances for the repeats HERVL (D), CR1_Mam (E), and OldhAT1 (F) in the human genome, using both HMMER (filtered using the GA threshold provided in the Dfam models) and BLAST (filtering using e-value <=1e-05).

To determine how the increased effectiveness of HMMER over BLAST varies with repeat age, we stratified repeat models from the Dfam database according to the estimated timing of transposition and calculated the fold-change in bases detected by HMMER relative to BLAST in the *H. sapiens* genome (Fig. 1B). As expected, very young, human-specific repeat models were detected in nearly identical amounts by both methods, consistent with the limited sequence divergence accumulated since insertion and the fact that BLAST’s seed-based search approach remains equally effective to HMMER’s in the absence of sequence decay. In contrast, progressively older groups of repeats showed a marked increase in HMMER sensitivity: models assigned to the Primate, Euarchontoglires, Boreoeutheria, and Eutheria ancestors exhibited median gains of approximately 1.61-fold, 1.87-fold, 1.84-fold, and 3.12-fold, respectively, in coverage relative to BLAST. This trend became even more pronounced for ancient repeats, with models dating to the Theria, Mammalia, Amniota, and Tetrapoda ancestors showing median gains of 8.11-fold, 6.50-fold, 4.77-fold, and 4.64-fold, respectively. Notably, HMMER outperformed BLAST for 100% of repeat models analyzed in the *H. sapiens* genome and for 99.2% of models in the *M. musculus* genome (Fig. 1C). Together, these results demonstrate the substantially greater sensitivity of HMMER for the detection of ancient repeats and indicate that reconstruction approaches targeting these ancient TEs would benefit from HMMER-based searches.

To understand how the difference in sensitivity between BLAST and HMMER affects the extension process for highly degenerate repeat families, we examined how coverage varied along the length of three repeat models of different evolutionary ages: HERVL (Euarchontoglires), CR1_Mam (Mammalia), and OldhAT1 (Tetrapoda). These repeat families span a range of fragmentation and divergence levels and are therefore informative of the challenges encountered by repeat extension algorithms. In the case of HERVL, BLAST and HMMER detected a similar number of insertions, and despite substantial fragmentation, many insertions remained full-length or nearly full-length, achieving relatively uniform coverage along the model (Fig. 1D). In contrast, for the more ancient and degraded CR1_Mam and OldhAT1 families, HMMER yielded markedly greater coverage than BLAST, with 10-fold and 4.8-fold increases in overall coverage, respectively (Fig. 1E,F). This improvement was accompanied by increases in both the number and length of detected instances: the number of insertions rose by 266.0% for CR1_Mam and 136.89% for OldhAT1, while median insertion length increased from 172 bp to 321 bp for CR1_Mam and from 168 bp to 269 bp for OldhAT1. Strikingly, some regions of the repeat models were so highly decayed that they could only be detected with HMMER, indicating that their reconstruction is unlikely to be possible using BLAST. Additionally, these ancient elements were also extremely fragmented, and no full-length insertions were recovered, even with HMMER. In the case of OldhAT1, five regions of elevated conservation could be noted along the repeat model, yet individual fragments rarely spanned across more than two of them. This pattern poses a major challenge for existing repeat reconstruction approaches, since even when extension is initiated in any one of these conserved regions, current algorithms are likely to recover only the immediately neighboring region and fail to extend into the next, owing to the absence of fragments spanning all three. Thus, whereas BLAST-based extension strategies that retrieve immediately adjacent genomic sequence may be sufficient for the reconstruction of relatively recent repeats, the reconstruction of ancient TEs is likely to require HMM-based search strategies together with dynamic, iterative procedures capable of linking fragments connected only through transitive adjacencies.

### RRE uses a dynamic search strategy to improve repeat models

To overcome the limitations of repeat extension approaches based on the BEEA paradigm, we developed Recursive Repeat Extender (RRE), a Nextflow-based (DI Tommaso et al., 2017) computational pipeline for automated repeat library extension. Whereas BEEA employs a static search strategy, in which repeat instances are identified once and then extended (Fig. 2A-C), RRE uses a dynamic search approach in which the repeat model extended in one round is used to query the genome again in the next (Fig. 2D-F). This recursive procedure enables the progressive merging of repeat regions linked only through transitive adjacencies. In addition, RRE uses HMMER rather than BLAST for repeat detection, allowing the reconstruction of highly degenerated repeats.

**Figure 2.**
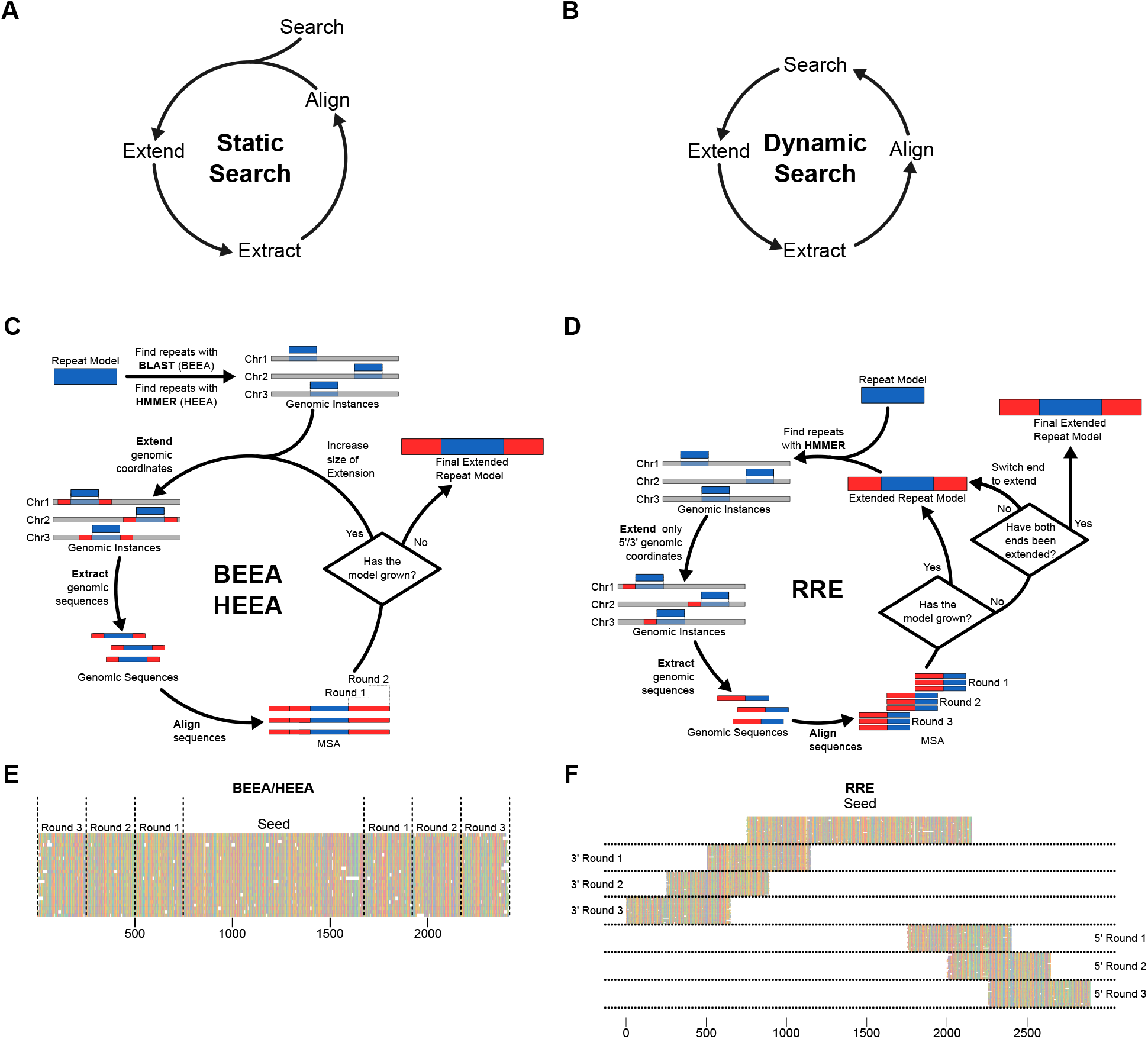
Differences between BEEA and RRE algorithms. A-B) Schematics showing the general algorithm of the static search (A) and the dynamic search (B) on which BEEA/HEEA and RRE are based, respectively. C) Schematic illustrating the BEEA/HEEA algorithms: after an initial search using BLAST (BEEA) or HMMER (HEEA), these algorithms proceed to extend the genomic coordinates of the repeats found and then extract and align the corresponding genomic sequences. After alignment, an assessment is made to determine whether the model has grown; if it has, the algorithm extends the model further; otherwise, the extension stops. D) Schematic illustrating the RRE algorithm, which starts by searching repeat models using HMMER. Unlike BEEA, RRE extends one end at a time (either 5’ or 3’). The genomic sequences corresponding to the extended coordinates are then extracted and aligned. Once aligned, the aligned sequences are joined to the model from the previous round using MAFFT’s --add functionality. Then, the algorithm determines whether the model has grown; if it has, it launches a new round of extension using the updated version of the model as the search query. Once extension of one stops yielding increases in length, the algorithm moves on to the other end. Once both ends have been extended, the algorithm finalizes and moves on to the model polishing step (Module 4). E) and F) Example alignments produced by BEEA/HEEA (E) and RRE (F) algorithms after 3 rounds of extension. Each horizontal line represents an individual genomic locus. BEEA/HEEA generate extensions relative to an initial search step and therefore the employed loci are fixed from the beginning of the algorithm. In contrast, RRE repeats the search step in each round, incorporating new loci as necessary.

RRE requires a repeat library as input that contains repeat models in both Stockholm (.stk) and FASTA (.fa) formats, such as those generated by RepeatModeler2 (Flynn et al., 2020). RRE comprises five modules (described in detail in Suppl. Methods, Fig. 3):

**Figure 3.**
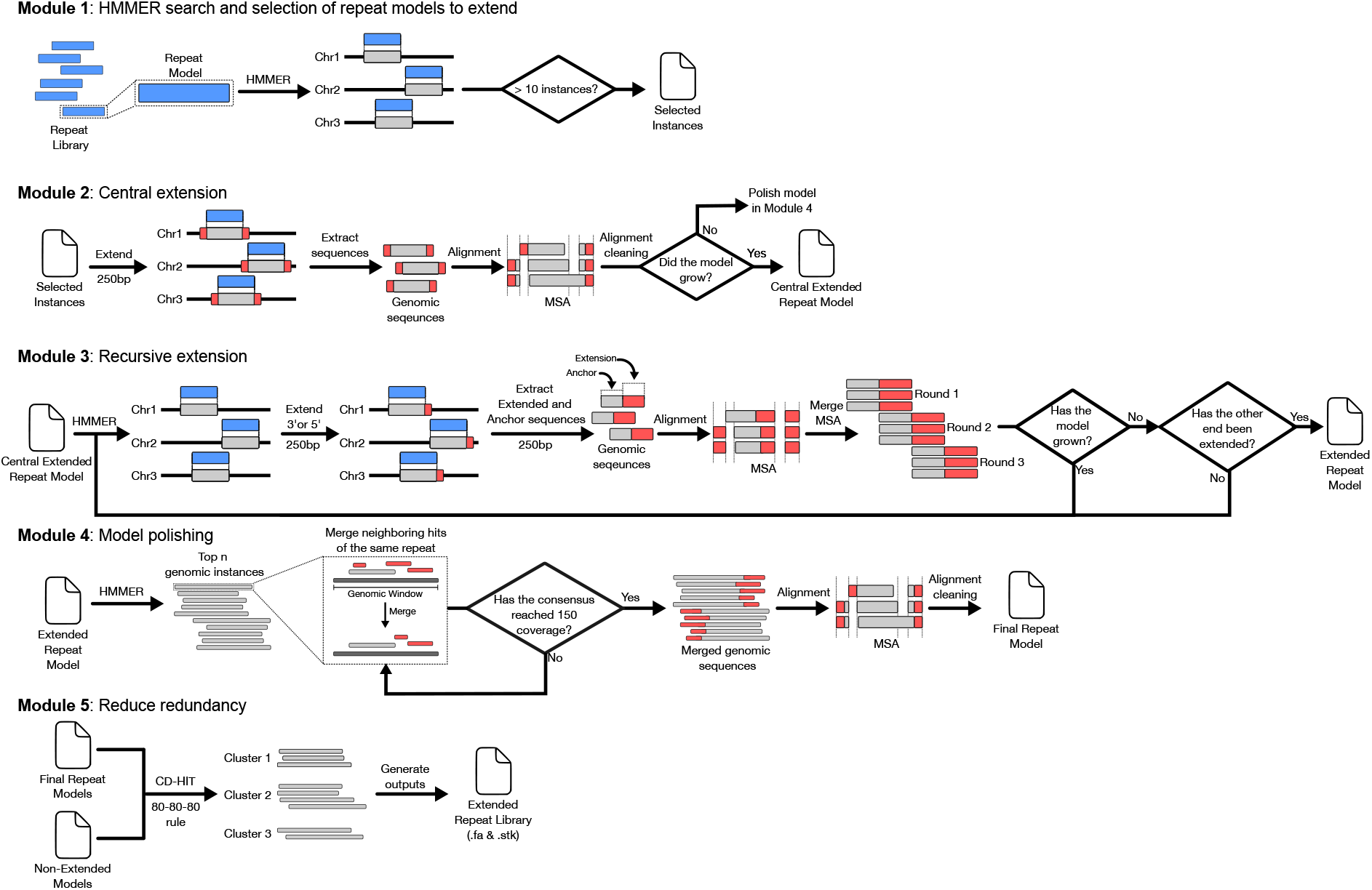
REE workflow to extend repeats. Red denotes extended sequences, grey denotes genomic sequences, and blue denotes repeat models. The pipeline is organized into five sequential modules. Module 1 performs the initial search with HMMER and selects repeat models with more than 10 genomic instances, which are then extended in subsequent steps. Module 2 performs the central extension by extending the genomic coordinates found in the initial search in Module 1 by 250 bp bidirectionally. The genomic sequences from the extended genomic coordinates are then extracted and aligned. After the alignment has been cleaned, the algorithm determines whether the model has grown and from which end; if it has grown, regardless of the end, the repeat model is extended further in Module 3; otherwise, the model skips Module 3 and is used directly in Module 4. Module 3 then performs the recursive extension by searching the repeat model using HMMER at the beginning of each round and selecting only those instances that contain sequences corresponding to the end being extended. Next, the algorithm extends the genomic coordinates by 250bp in one direction only (the 5’ end is done first, then the 3’). The genomic sequences corresponding to the extended coordinates are then extracted and aligned. Once the alignment is cleaned, the algorithm merges it with the alignments from previous rounds. At the end of the cycle, the algorithm determines if the model has grown; if it has, it proceeds to do another round of extension; if not, then the algorithm determines if it proceeds to extend the remaining side (3’ end) or stop the extension. Module 4 performs model polishing by searching for the extended repeat model in the genome using HMMER and ordering the instances by bit score. Iterating through the instance list, this module searches the genomic neighborhood of each instance and joins smaller fragments within a specified window. The module iterates through the instance list until it has enough instances to achieve 150 bp coverage at each base. The module subsequently aligns the genomic sequences corresponding to the merged instances and produces a final repeat model. Lastly, Module 5 produces a final repeat by library by merging the extended and non-extended models. To reduce redundancy, this model clusters the models using CD-HIT and chooses which model to keep based on the 80-80-80 rule (80% identity, 80% coverage across at least 80 bp).

#### Module 1: HMMER search and selection of repeat models to extend

This module generates HMM models from the Stockholm alignments and uses HMMER to search the genome for each repeat model. Then, this module determines whether each repeat model has sufficient genomic instances to be extended, using an arbitrary threshold (a minimum of 10 instances is required by default).

#### Module 2: Central extension

This module performs an initial extension by extending each instance identified in Module 1 by a fixed length (e.g., 250 bp) at both ends, followed by a MSA of the resulting genomic sequences. It then determines whether either end of the repeat model has increased in length; if so, the expanded ends are extended further in Module 3.

#### Module 3: Recursive extension

This module extends the ends of the repeat model if they extended successfully in the previous module. If both ends are to be extended, they are extended sequentially (the 5’ end is extended first). This is done to avoid merging the alignments produced by both ends and to avoid conflicts with models generated by our family-splitting approach (see Suppl. Methods). Each round of extension starts by searching the complete repeat model from the previous round (or, in the first round, the extended model from Module 2), and it retains only those search hits that overlap with the previous round’s extended region. Using these repeat instances, this module extends the genomic coordinates by a set amount (250 bp by default) towards one direction (3’or 5’) in reference to the repeat model orientation. Then the module uses the genomic sequences corresponding to the extended instances to generate an MSA. The resulting MSA is then merged with the MSAs from the previous round using MAFFT’s “`--add`” command. If the model successfully grows in length, another round of extension is undertaken. Once the model no longer shows an increase in length, the extension process terminates, and the module either moves on to the other end (if Module 2 determined it can be extended) or moves on to Module 4 (once both ends have been addressed).

#### Module 4: Model polishing

This module searches the genome using the final model generated by module 3 using HMMER, constructing a new repeat model that prioritizes the most contiguous repeat instances and incorporates a larger number of sequences (aiming for a coverage of 150 at every base of the consensus by default). Repeat instances in the same genomic neighborhood (10 kbp by default) are concatenated and handled as a single fragment to account for repeats that might have been disrupted by an insertion event. This module is necessary to mitigate errors that could have been introduced by the recursive search approach (e.g. resolve ambiguities regarding the number and relative placement of repetitive segments such as LTRs) and to produce a more accurate model built on more sequences.

#### Module 5: Assembly of final repeat library

This module reduces redundancy in the repeat library, which comprises all extended repeat models and the repeat models not selected for extension by Module 1. The consensus sequences representing the repeat models in the library are clustered using CD-HIT (Fu et al., 2012), following the 80-80-80 rule (80% identity, 80% coverage, and 80 bp) (Goubert et al., 2022). The resulting repeat library is then output in FASTA and Stockholm format.

Additionally, RRE implements novel approaches/features that are used across several of its modules:

#### Adaptive MSA cleaning

Since ancient TEs are highly degenerate and often approach the twilight zone of sequence alignability, we implemented a novel adaptive method for MSA cleaning that adapts to the degree of sequence divergence. Instead of using conventional sequence identity as the criterion to determine whether a position in the MSA should be pruned, we adapted the trident estimator from Nguyen (2011), which accounts for information content and empty information at each position (see Suppl. Methods). This method, based on the distribution of trident estimator values, identifies which values are more likely to contain sequences not part of the repeat and generates an MSA that removes positions below that threshold

#### Family detection and splitting

In order to avoid chimeric models between different families containing shared conserved regions, at each round we undertake a family splitting step. In brief, this approach takes the MSA as input, computes the distribution of pairwise sequence identities across the alignment, and determines whether the distribution is multimodal. If it is found to be multimodal, it constructs a dendrogram via hierarchical clustering, splits the sequences into multiple groups based on the distribution of pairwise distances, and generates an MSA for each family (see Suppl. Methods). The extension pipeline subsequently branches, allowing each family to be extended independently in parallel

#### Comprehensive repeat search

Modules 2 and 3 search the entirety of the repeat model available until that point, as opposed to just the extended section from the previous round. This results in the pipeline prioritizing fragments that are part of larger repeat instances, thereby minimizing the risk of creating chimeric repeat models (see Suppl. Methods)

### RRE improves *de novo* repeat libraries

To assess the performance of the RRE approach, we used five genomes with extensively curated repeat annotations available in Dfam (*C. elegans, D. melanogaster, D. rerio, M. musculus*, and *H. sapiens*), as well as *de novo* annotations for each genome generated with RepeatModeler2. To compare the recursive approach with conventional methods, we implemented a version of the BEEA algorithm that uses HMMER instead of BLAST as the search algorithm, which we term HEEA (HMMER-Extend-Extract-Align). Comparing RRE with HEEA allowed us to control for the search method and isolate differences attributable specifically to the extension strategy of the two algorithms.

To assess the completeness of each repeat library, we followed the benchmark established by Flynn et al. (2020). In their benchmark, they compare the two libraries (reference and query) using RepeatMasker and assign models in the reference library a category depending on their percentage of identity and coverage in the query library. These categories are “Perfect” (models with 95% identity and 95% reciprocal coverage), “Good” (models with 95% identity and 95% coverage of the reference model over multiple models in the query library), “Present” (models with 80% identity and 80% coverage of the reference model over multiple models in the query library), and “Missing” (those models that did not meet any of the beforementioned criteria). To this framework, we made two modifications in order to benchmark extended repeat libraries: the first was the removal of the “Perfect” category, as the RRE and HEEA approaches don’t separate long terminal repeats (LTRs) from the LTR class elements, which would make these libraries incomparable to RepeatModeler2 and Dfam libraries, which separate the LTRs from the main repeat, and would lead to an underestimation of “Perfect” repeats in RRE and HEEA libraries. The second modification introduces the “Poor” category (>80% identity and >50% coverage across multiple repeat consensus sequences), which more accurately represents repeats that are present but fragmented in the reference library, such as the partial L1 segments that constitute the mammalian Dfam annotation. We note that given these substantial differences in library structure and repeat model representation, there is no one-to-one correspondence between HEEA/RRE-derived query models and Dfam reference models. Therefore, this benchmark should only be interpreted as a measure of relative performance.

Both RRE and HEEA had more “Good” repeat models than the repeat library generated by RepeatModeler2, with RRE having more “Good” repeat models than HEEA in *C. elegans, D. melanogaster*, and *D. rerio*, while HEEA had more “Good” models in *M. musculus and H. sapiens* (Fig. 4A). In detail, HEEA increased the number of “Good” repeat models by 1.5% in *C. elegans*, 6.8% in *D. melanogaster*, 5.2% in *D. rerio*, 3.6% in *M. musculus*, and 5.8% in *H. sapiens*. By comparison, RRE increased the number of “Good” repeat models by 3.0% in *C. elegans*, 10.0% in *D. melanogaster*, 8.7% in *D. rerio*, 2.8% in *M. musculus*, and 2.3% in *H. sapiens*. In summary, HEEA and RRE increased the combined number of “Good” repeats, with RRE outperforming HEEA in *C. elegans, D. melanogaster*, and *D. rerio*, but HEEA outperforming RRE in *M. musculus* and *H. sapiens*.

**Figure 4.**
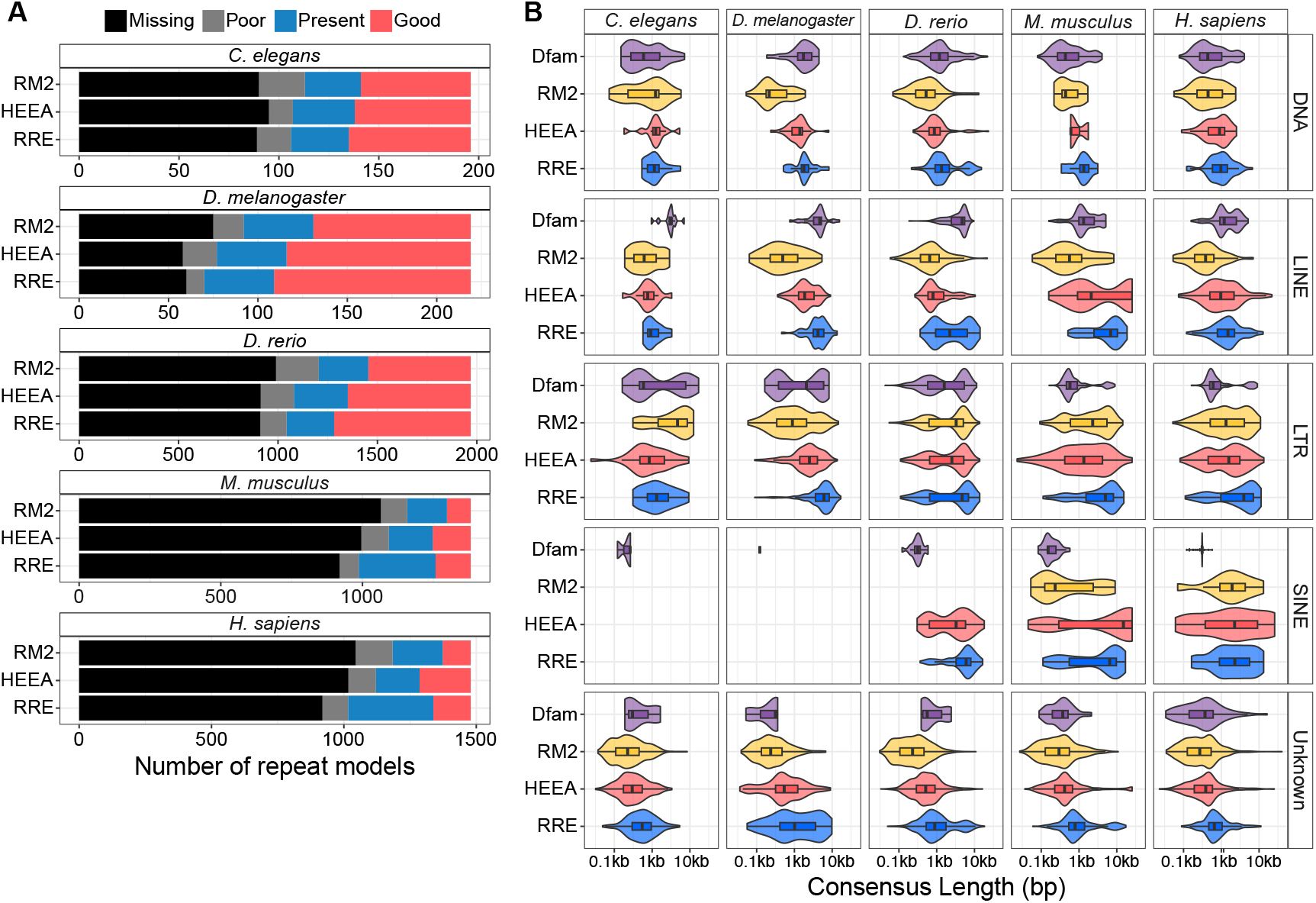
Benchmarking RRE performance extending repeat annotations. A) Consensus classification based on Flynn et al. (2020). “Good” are reference repeat models covered across multiple models in the subject library, with an identity >95% and a coverage >95%. “Present” refers to reference models covered across multiple models in the subject library, with identity >80% and coverage >80%. “Poor” are reference models covered across multiple models in the subject library, with an identity >80% and a coverage >50%. “Missing” are reference repeat models that couldn’t be classified in the previous categories. B) Repeat model length distribution of the main repeat classes (DNA, LINE, LTR, SINE, and Unknown) in five different species.

Another important goal of extension approaches is to reduce the number of “Missing” and “Poor” repeats. Compared to the repeat library generated by RepeatModeler2, HEEA and RRE reduced the number of “Missing” and “Poor” entries after extension. HEEA reduced the number of “Missing” repeat models by 7.7% in *D. melanogaster*, 3.9% in *D. rerio*, 4.9% in *M. musculus*, and 1.8% in *H. sapiens*, but increased by 2.5% in *C. elegans*; while RREA reduced this number by 0.5% in *C. elegans*, 6.8% in *D. melanogaster*, 4.0% in *D. rerio*, 10.5% in *M. musculus*, and 1.8% in *H. sapiens*. “Poor” annotations were also reduced by both extension approaches, with HEEA reducing the number of these annotations by 5.6% in *C. elegans*, 2.4% in *D. rerio*, and 2.4% in *H. sapiens*, but increasing by 0.9% in *D. melanogaster* and 0.2% in *M. musculus*. By contrast, RRE reduced the number of “Poor” annotations by 3.0% in *C. elegans*, 3.2% in *D. melanogaster*, 4.2% in *D. rerio*, 1.8% in *M. musculus*, and 2.9% in *H. sapiens*. Taken together, both extension approaches reduced the number of “Missing” and “Poor” repeat models, with RRE showing the greatest improvement (Fig. 4A).

### RRE increases the length of repeat models

*De novo* repeat libraries often contain shorter repeat models compared to manually curated libraries (Suppl. Fig. 1). After extension, repeat models from all major repeat classes are expected to be longer, ideally approaching the length of the models present in reference libraries. In general, we observed an increase in the length of repeat models in the most abundant repeat classes (DNA, LINE, LTR, SINE, and Unknown) after extension through the RRE and HEEA algorithms (Fig. 4B). RRE annotations tend to be longer than those extended using HEEA, observing overall median increases of 81.4% in *C. elegans*, 226.1% in *D. melanogaster*, 78.8% in *D. rerio*, 152.4% in *M. musculus*, and 91.6% in *H. sapiens*.

RRE approached the biological expected lengths for some of the repeat classes (Fig. 4B). For example, RRE generated models for LINE elements whose median length is longer than 3000 bp in some species (*M. musculus* and *D. melanogaster*), whereas the RM2 and HEEA libraries contain LINE models with a median length under 2000 bp (Fig. 4B). For LTR repeats, RRE-generated libraries have a median length over 5000 bp, while HEEA libraries have a median of under 3000bp for all species (Fig. 4B). Finally, in repeat libraries generated by RRE, the median length of Unknown repeat models was more than twice that in *de novo* libraries, but these models remain unannotated due to the absence of protein-domain features and likely represent non-autonomous elements (Fig. 4B). Overall, RRE was able to increase the length of repeat models across all species and classes.

### RRE reduces the size of repeat libraries and improves overall repeat identification

Repeat extension approaches aim to reduce the number of repeat models by increasing the length of each repeat model and discarding models with redundant sequences. After extension, HEAA reduced the number of consensus by 8.9% in *C. elegans*, 23.4% in *D. melanogaster*, 19.3% in *D. rerio*, 19.7% in *M. musculus*, and 9.5% in *H. sapiens*; while RRE reduced the library size by 18.7% in *C. elegans*, 28.1% in *D. rerio*, 6.1% in *M. musculus*, and 15.0% in *H. sapiens*. However, RRE resulted in an increase of 14.4% in *D. melanogaster* due to a high amount of LTR repeats (Fig. 5A; see Discussion). Overall, we observed a decrease in the number of repeat consensuses after extending with HEEA and RRE, with RRE producing the libraries with the lowest number of repeat models.

**Figure 5.**
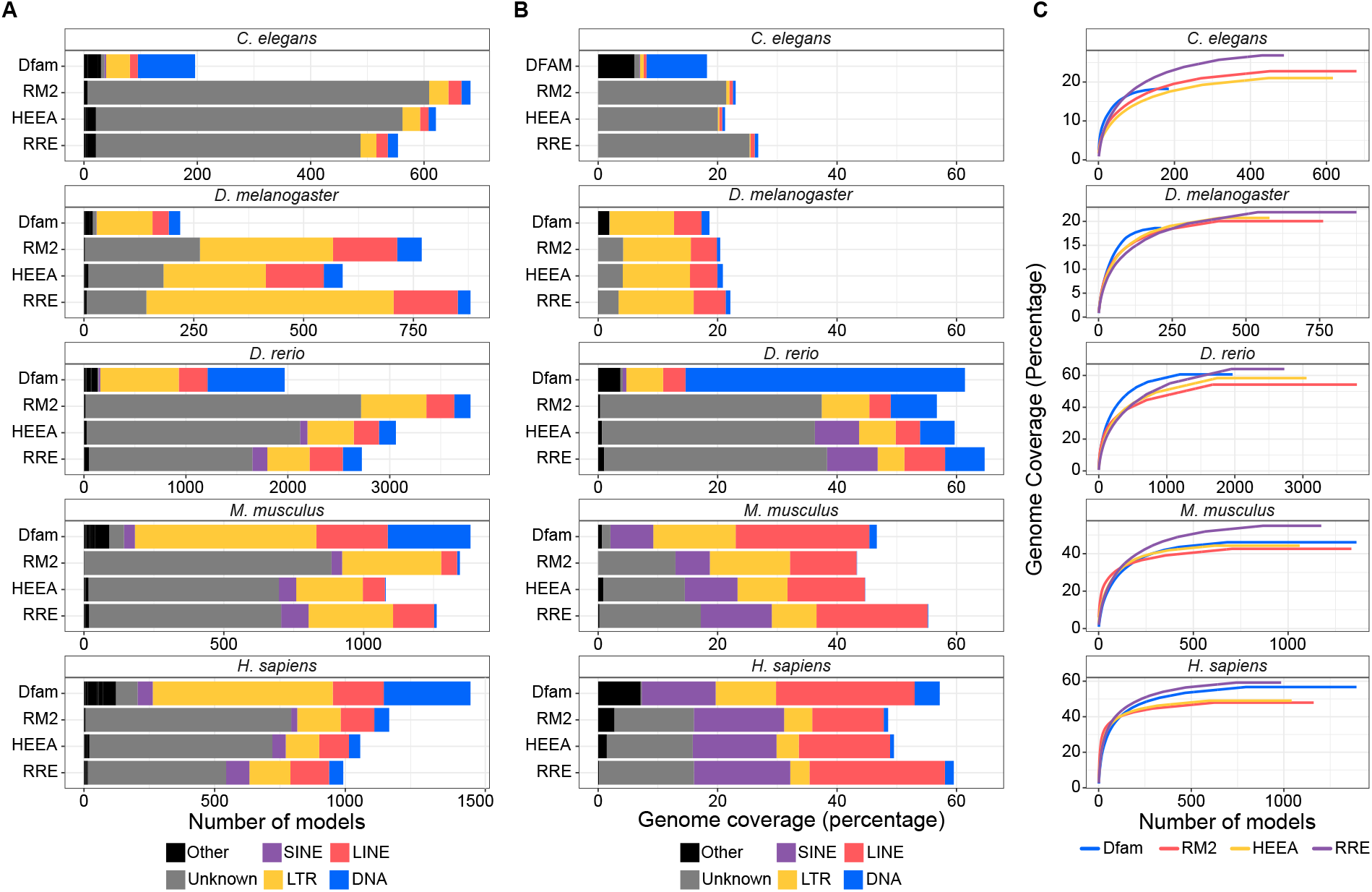
RRE improves repeat libraries across multiple species. A) Number of consensuses present in four different repeat libraries (Dfam, RM2, HEEA, and RRE) across five different species. Colors denote the repeat classification, with the Others class representing repeats that are not classified in the other depicted classes (e.g., Satellites, tRNAs). B) Genome coverage achieved after masking the genome with four different repeat libraries (Dfam, RM2, HEEA, and RRE) across five different species. Colors denote the repeat classification, with the Others class representing repeats that are not classified in the other depicted classes (e.g., Satellites, tRNAs). C) Saturation curves showing the cumulative sum of genomic coverage using three different repeat annotations (Dfam, RM2, HEEA, and RRE).

Repeat libraries generated with RepeatModeler2 contain numerous unclassified repeat models, many of which are likely fragments of longer repeats and non-autonomous repeats lacking protein domains (e.g., MITEs, SINEs and non-autonomous LTRs). While the number of repeat models corresponding to the main repeat classes remained mostly unchanged, extended repeat libraries with HEEA and RRE have fewer models classified as Unknown (Fig. 5A). With HEEA, Unknown models were reduced by 10.2% in *C. elegans*, 34.7% in *D. melanogaster*, 22.5% in *D. rerio*, 22.9% in *M. musculus*, and 11.0% in *H. sapiens* compared to the *de novo* annotation. In contrast, RRE reduced Unknown repeat models by 22.5% in *C. elegans*, 48.0% in *D. melanogaster*, 40.6% in *D. rerio*, 22.2% in *M. musculus*, and 32.7% in *H. sapiens*. In general, RRE outperformed HEEA in reducing the number of repeats classified as Unknown, resulting in fewer models in the repeat library.

To further assess the quality of the extended repeat libraries, we used them to identify repeat content in their respective genomes. In general, both extension methods generated libraries that identified more of the genome as repeats compared to libraries generated by RepeatModeler2 (Fig. 5B). The repeat library generated by the HEEA algorithm increased the total fraction of the genome identified as repeats by 0.4% in *D. melanogaster*, 2.97% in *D. rerio*, 1.3% in *M. musculus*, and 0.9% in *H. sapiens* of the overall genome. In contrast, RRE increased the percentage of the genome identified as repeats by 3.7% in *C. elegans*, 1.7% in *D. melanogaster*, 8.0% in *D. rerio*, 11.9% in *M. musculus*, and 10.9% in *H. sapiens*. In conclusion, the repeat libraries generated by RRE outperformed the libraries from HEEA in identifying the repetitive fraction of the genome.

Lastly, we examined saturation curves for each repeat library (Dfam, RepeatModeler2, HEEA, and RRE) to assess how rapidly each library increased the fraction of the genome annotated as repetitive (Fig. 5C). RepeatModeler2 libraries showed the steepest initial rise, exceeding even Dfam, which we attribute to the generation of chimeric models derived from highly abundant repeats. Among the extension-based libraries, RRE exhibited a steeper accumulation curve than HEEA, indicating that it annotated more repetitive sequences using fewer models. In contrast, HEEA libraries reached saturation earlier and plateaued at lower levels than RRE libraries. Overall, these results indicate that RRE generates more comprehensive and less redundant repeat libraries, enabling broader genome annotation before reaching saturation.

### REE can reconstruct ancient TEs

Remnants of ancient TEs dating back hundreds of millions of years are scattered throughout vertebrate genomes. These elements are often highly fragmented and exceptionally divergent, making their reconstruction particularly challenging. To our knowledge, no publicly available pipeline capable of reconstructing ancient TEs is currently available. To enable the reconstruction of such highly degraded elements, we developed an alternative RRE mode, termed “AncientMode”, specifically optimized for ancient repeats. AncientMode includes an alignment-cleaning strategy tailored to highly degenerate sequences, omits repeat-family detection steps, builds HMMs only from the fragment currently being extended in order to maximize recovery of insertions overlapping the distal edge, and applies a more stringent procedure for merging MSAs across successive rounds of extension (see Suppl. Methods).

To test whether RRE’s AncientMode can successfully recover ancient TEs, we sought to reconstruct the mammalian CR1 repeat, CR1_Mam, which is annotated in Dfam and inferred to have been active in a common mammalian ancestor approximately 180 million years ago (Kumar et al., 2022). To mimic a realistic *de novo* discovery scenario, we began with a truncated seed model containing only the most abundant portion of the repeat, corresponding to positions 1476–2204 of the Dfam consensus (Fig. 6A). Using RRE in AncientMode, we recovered fragments spanning the full CR1_Mam repeat and extended the reconstructed model 131 bp beyond the boundaries of the curated Dfam consensus after 22 rounds of extension (Fig. 6B). We then calculated gathering (GA) thresholds for both the RRE-derived model and the Dfam model using our RepeatMasker-based pipeline (see Suppl. Methods) and searched the human genome with each model. The RRE-derived model achieved genomic coverage comparable to that of the curated Dfam model (Fig. 6C-D). In addition, the individual repeat insertions detected by the two models were broadly similar in size distribution (Fig. 6E-F), with median insertion lengths of 213 bp for the Dfam model and 255 bp for the RRE-derived model. Overall, these results show that RRE can reconstruct ancient repeats from a seed alignment, achieving genomic coverage comparable to that obtained with a curated Dfam model while also recovering previously unannotated regions beyond the current model boundaries.

**Figure 6.**
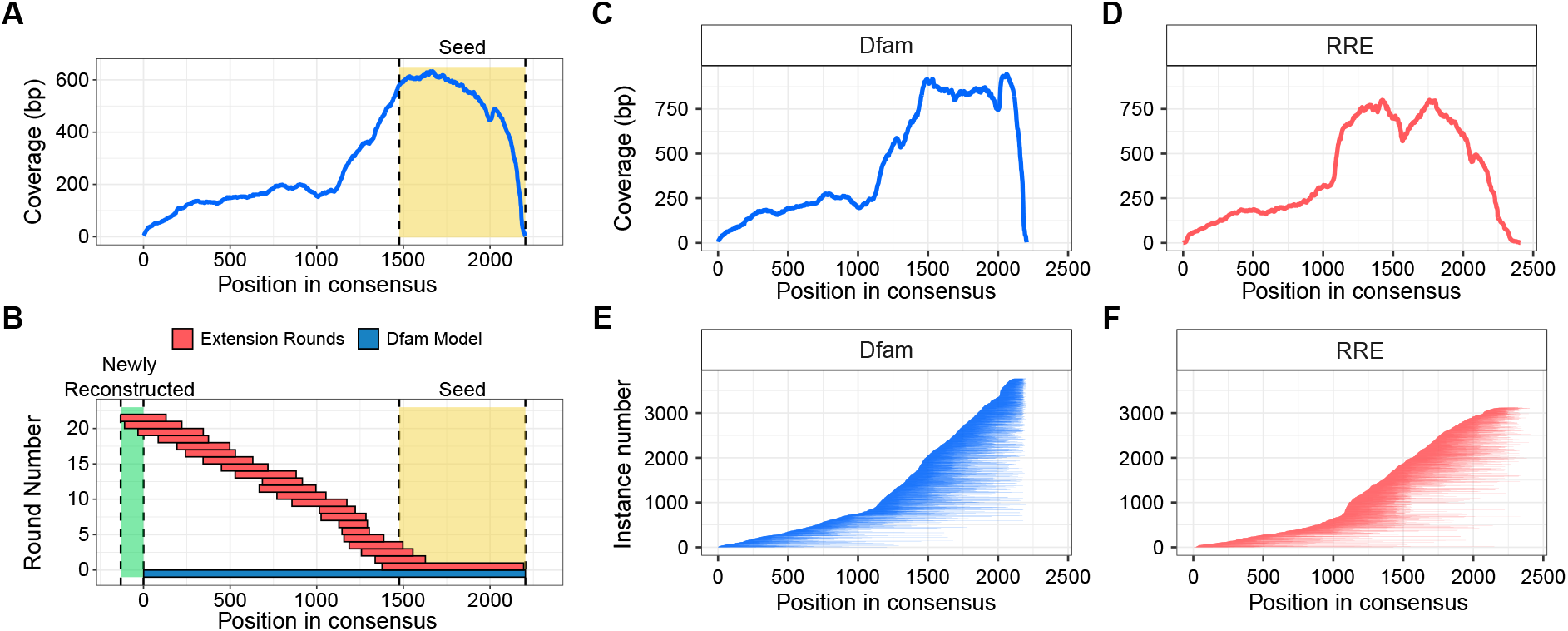
RRE reconstructs the CR1_Mam sequence. A) Schematic showing the coverage of instances found using the Dfam curated model with HMMER (using the GA threshold provided by Dfam as the threshold), and the highlighted region represents the section of the model used for the generation of the truncated repeat model. B) Schematic showing the regions covered by each round of the extension (red) relative to the Dfam model (blue). The yellow highlighted region represents the truncated model that was used as seed model, while the green highlighted region represents the newly reconstructed region that was missing from the curated Dfam model. C) and D) Coverage of the repeat model using the instances in the human genome found by the curated Dfam CR1_Mam model (C) and the RRE-extended CR1_Mam repeat model (D). E) and F) Individual instances of CR1_Mam found using the curated Dfam model (E) and the RRE-extended repeat model (F), ordered by starting position in their respective consensus.

## Discussion

We introduce RRE, a new approach for improving repeat libraries that overcomes two major limitations of conventional repeat-extension methods: their reliance on seed-based repeat searches and their dependence on a single genomic search to provide the substrates for extension. We addressed the first limitation by adopting HMMER as the primary engine for repeat detection. Because profile HMMs use position-specific probabilistic models, HMMER is substantially more sensitive than BLAST for the detection of highly degenerate repeats (Fig. 1A), and this advantage becomes especially pronounced for repeat families that were active around the emergence of the Mammalia, Amniota, and Tetrapoda clades (Fig. 1C). We addressed the second limitation through the recursive design of RRE, in which iterative cycles of search and extension progressively incorporate fragments distributed across multiple genomic loci. This strategy maximizes the amount of information recovered from the genome and enables the reconstruction of repeat families that no longer retain intact copies (Fig. 3). Together, these advances allow RRE to generate improved repeat libraries within a scalable and reproducible framework implemented in Nextflow.

We used RRE and an implementation of the BEEA algorithm that employs HMMER instead of BLAST (HEEA) to extend *de novo* repeat libraries for five species (*C. elegans, D. melanogaster, D. rerio, M. musculus*, and *H. sapiens*). RRE reduced the number of missing annotations and improved the quality of existing ones (Fig. 4A). However, the interpretation of such comparisons remains limited by the lack of standardized benchmarks for repeat annotation methods and by the heterogeneous ways in which repeat families are represented across methods and reference libraries. For example, some libraries separate LTRs from internal LTR-retrotransposon regions or split LINEs into multiple component models, complicating direct comparisons. More standardized benchmarking frameworks and reference libraries will therefore be necessary to evaluate the accuracy and performance of repeat annotation tools more rigorously (Hoen et al., 2015). The greater completeness of RRE-extended libraries is also reflected in repeat model length, as the models generated by RRE were generally longer than those produced by RepeatModeler2 and HEEA (Fig. 4B). These results indicate that RRE yields longer and more complete repeat models than both the original d*e novo* libraries or libraries extended using HEEA.

A major goal of repeat-extension approaches is to reduce both the total number of repeat models and the number of elements classified as Unknown, as extension can merge fragmented or partial models into longer consensus sequences that are more readily classified. In general, both extension methods reduced the number of repeat models relative to the *de novo* libraries (Fig. 5A). RRE further outperformed HEEA in reducing library size in all species except *D. melanogaster*, where we observed an increase specifically in the number of LTR element models, but not in other repeat classes (Fig. 5A). We hypothesize that this pattern reflects inter-element recombination among LTR retrotransposons, a phenomenon previously described in *D. melanogaster* (Jordan & McDonald, 1998; Senti et al., 2025; Vargiu et al., 2016). Such recombination may generate mosaic or divergent structures that are subsequently split into multiple families by our family splitting procedure (see Suppl. Methods).

To assess the effectiveness of the resulting repeat libraries, we used RepeatMasker to annotate repeats in their respective genomes. Libraries produced by both extension approaches identified a larger fraction of the genome as repetitive than the original *de novo* libraries (Fig. 5B). However, only RRE increased the fraction of the genome annotated as repetitive beyond that identified using the Dfam reference library (Fig. 5B).

We also provide a proof of principle that RRE can reconstruct ancient repeats from a truncated seed model. RRE was able to reconstruct the CR1_Mam repeat from a truncated seed despite the extensive fragmentation observed among its genomic instances in the human genome (Fig. 1E), and it further improved the model by extending it beyond the boundaries of the reference model available in Dfam (Fig. 6B). The resulting model achieved genomic coverage comparable to that of the curated Dfam model (Fig. 6C), while detecting longer individual fragments (Fig. 6D). These results demonstrate that RRE can reconstruct ancient repeats from fragmentary seed models. Further improvements may come from incorporating alignment methods specifically tailored to highly degenerate sequences, such as Refiner (Hubley et al., 2022). However, to be compatible with the recursive search strategy implemented in RRE, such methods would need to support the incorporation of newly identified sequences into pre-existing alignments across successive extension rounds, similarly to MAFFT’s “`--add`” functionality.

RRE is intended to enable the annotation of ancient repeats, which have been shown to be sources of enhancers (Bejerano et al., 2006; Lynch et al., 2011) and other *cis*-regulatory elements (Wenger et al., 2013). By generating improved models of ancient repeats, we aim to better understand their origins and to support their identification across related species. With continued advances in whole-genome alignment methods that improve the identification of conserved noncoding regions (Armstrong et al., 2020), we expect RRE to become a useful tool for annotating ancient repeats in both extant genomes and ancestral genome reconstructions (Matsushima et al., 2024).

Overall, RRE provides an effective framework for automated repeat-library extension, outperforming conventional approaches and being specifically designed to enable the reconstruction of particularly fragmented or diverged repeat families. Its improved performance stems from the greater sensitivity of HMMER-based searches and the recursive extension strategy, in which the extended model from each round is used to seed the next. Because RRE is implemented in Nextflow and distributed with a Docker container, it is readily scalable and easily deployable in most high-performance computing (HPC) environments. Moreover, to our knowledge, RRE provides the first repeat-extension framework specifically tailored to ancient repeats, and our CR1_Mam reconstruction demonstrates its ability to recover such elements in practice.

## Methods

### Genomes used for repeat annotation and extension

We used the *Caenorhabditis elegans* WBcel235 assembly (GCA_000002985.3), *Drosophila melanogaster* Release 6 plus ISO1 MT assembly (GCA_000001215.4), *Danio rerio* GRCz11 assembly (GCA_000002035.4), *Mus musculus* GRCm39 (GCF_000001635.27), and *Homo sapiens* T2T-CHM13v2.0 assembly (GCF_009914755.1) from the NCBI database for all *de novo* annotations and repeat extensions.

### Reference annotations

Repeat annotations for *Caenorhabditis elegans, Drosophila melanogaster, Danio rerio, Mus musculus, and Homo sapiens* were obtained from Dfam version 3.8 in FASTA and Stockholm formats. For individual searches, we used the Dfam repeat models for HERVL (DF000000198), Mam_CR1 (DF000000110), and OldhAT1 (DF000001275) in FASTA and HMM formats.

### RepeatModeler2 for raw repeat libraries

All genomes listed above were used for repeat annotation using a modified version of RepeatModeler v2.0.6 (https://github.com/BioFalcon/RepeatModeler), which was modified to parallelize the repeat annotation by splitting the LTRPipeline from the RECON/RepeatScout modules. To run RepeatModeler2, we split the genome into 10Mb slices and generated the RepeatModeler2 index using the `BuildDatabase` command. We then used the RepeatModeler2 command, modifying the `RepeatScout` sample with the `-rsSampleSize` parameter to set the first-round sample size to 1.5% of each genome, and with the `-targetSamplingSize` parameter to ensure that RepeatModeler2 sampled at least 15% of each genome. For the LTR identification, we used the `LTRPipeline` script with default parameters. To merge the results, we used the `LTRMerge` script from the modified RepeatModeler2 repository.

### HMMER and BLAST

To compare the difference between HMMER and BLAST, we searched each Dfam library using BLAST (v.2.14.0), which was used with parameters `-word-size 7 -gapopen 2 -gapextend 2 - penalty -1 -reward 1`, and using a threshold of e-value of 1e-05; and HMMER (v3.4), using nhmmer with the `--cut_ga` parameter.

### HEEA extension

All HEEA extensions were done using the RRE pipeline (https://github.com/BioFalcon/RRE), using Nextflow V25.04.7, and the command `nextflow ./RRE.nf --Genome ./<GENOME>.fa -- outDir ./<OUTDIR> --consensus ./<CONSENSUS>.fa --extension 250 --hyperT -- consensusAln ./<CONSENSUS>.stk --Algorithm HEEA --noiseThreshold 0.20`, using the libraries generated by RepeatModeler2 as input.

### RRE extension

All HEEA extensions were done using the RRE pipeline (https://github.com/BioFalcon/RRE), using Nextflow V25.04.7, and the command `nextflow ./RRE.nf --Genome ./<GENOME>.fa -- outDir ./<OUTDIR> --consensus ./<CONSENSUS>.fa --extension 250 --hyperT -- consensusAln ./<CONSENSUS>.stk --Algorithm RRE --noiseThreshold 0.20`, using the libraries generated by RepeatModeler2 as input.

### Repeat library completeness assessment

The classification of repeat libraries was done using the script described in Flynn et al. (2020), making the comparison using RepeatMasker (v4.1.8) with the command `RepeatMasker -lib./<Dfam>.fa -nolow -pa 4 ./<CONSENSUS>.fa, and analyzing the output using the script get_family_summary_paper.sh` provided in https://github.com/jmf422/TE_annotation/

### RepeatMasker

The identification of repeats was done with the Nextflow pipeline provided in https://github.com/BioFalcon/RepeatMasker_HMMER using the following command line `nextflow./RepeatMasker_HMMER.nf --Genome <GENOME>.fa --consensus <CONSENSUS>.stk -- chunkSize 10000000 --outDir <OUTDIR> --chromSizes <GENOME>.fa.fai -- numModels 20`.

### CR1_Mam extension

Reconstruction of CR1_Mam was performed by first cropping the reference model in Stockholm/.stk format obtained from Dfam (DF000000110), then constructing the HMM model using `hmmbuild -- fragthresh 1 --hand CroppedCR1_Mam.hmm CroppedCR1_Mam.stk` and obtaining the consensus sequence using the command `hmmemit -c CroppedCR1_Mam.hmm`. Using the cropped Stockholm file, we ran the RRE Nextflow (V25.04.7) pipeline with the command nextflow ./RRE.nf --Genome ./Homo_sapiens.genome.fa --outDir ./<OUTDIR> --consensus ./CroppedCR1_Mam.fa --extension 90 --hyperT --consensusAln ./CroppedCR1_Mam.stk --Algorithm Recursive --SampleSeqs 150 -- prevRoundCoverage 0.25 --minSequenceRound 5 --AncientMode -- percentileHorizontal 0.3 --maxRounds 45. Due to a pair of regions sharing weak homology towards the end of the model, we could not obtain a repeat model incorporating all fragments in an automated manner. Instead, we obtained a model containing all fragments generated during the first 18 Rounds, and an additional model containing the last 3 rounds. To merge them, we used MAFFT’s `--seed` option to generate a complete repeat model.

## Data Availability

The repeat libraries reported in this study as well as the truncated and extended CR1_Mam model are available in Zenodo using the following identifier: 10.5281/zenodo.19579763

## Acknowledgements

This work was supported by funding from the Austrian Science Fund (FWF) (EMT, 10.55776/I4353) and from the ERC (EMT, AdG 742046 RegGeneMems). DRT acknowledges funding from EMBO (DRT, ALTF 907-2019), from the European Union’s Horizon 2020 program under the Marie Skłodowska-Curie grant agreement 101033244, and from the Austrian Science Fund (FWF) (DRT, 10.55776/PAT8372424). We would like to thank Oleg Simakov for his input during the development of this project and Kirsten Senti for his valuable comments on this manuscript. We would also like to acknowledge the technical assistance provided by the VBC HPC-IT team. The computational results presented here were obtained using computational resources from the CLIP cluster (https://clip.science) and the Austrian Scientific Computing (ASC) infrastructure (VSC5 and MUSICA).

## Author contributions

F.F. performed computational analyses and wrote computational pipelines. F.F., E.M.T. and D.R-T. conceptualized the algorithm. F.F., E.M.T. and D.R-T. wrote and revised the manuscript. E.M.T. and D.R-T. conceived the project, supervised the study and secured funding.

## Supplementary Figures

**Supplementary Figure 1.**
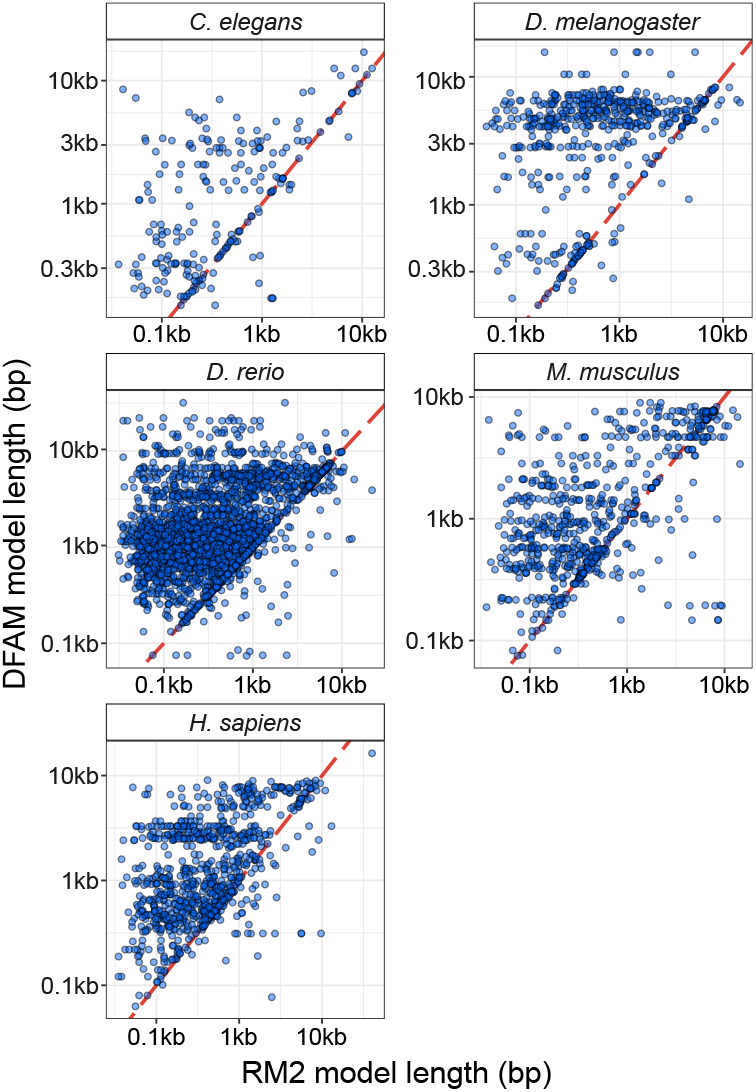
Comparison of repeat consensus lengths in repeat libraries generated *de novo* and in reference annotations available in Dfam. Pairs were determined by selecting the best hits from bidirectional BLAST between libraries generated with RepeatModeler2 and reference libraries from Dfam.

## Supplementary Methods

### Recursive Repeat Extender

The Recursive Repeat Extender algorithm is a tool for extending repeat annotations using consensus sequences (FASTA format) and their alignment files (Stockholm/.stk format). RRE comprises five modules that must be executed sequentially.

#### Module 1: HMMER search and selection of models to extend

This module performs the initial repeat search in a genome and determines whether a repeat can be extended. To begin, we create a HMMER database from the genome file using the `makehmmerdb` command using default parameters. In parallel, this module generates HMM profiles for all alignments in the Stockholm (.stk) format alignment file using the `hmmbuild` command, with the parameters `-- symfrac 0 --fragthresh 1.0 --wnone`. Afterward, the hmm profiles are split into individual files and searched in parallel for each repeat using the nhmmer command with default parameters, discarding hits below a user-defined threshold (1e-05 by default). After all models have been searched against the genome, we select only those with at least 10 instances and exclude LTRs from the LTR class, as we maintain the internal fraction for extension.

### Module 2: Central extension

This module’s task is to generate the ends that will subsequently be extended individually. This module begins by using the coordinates obtained from the previous nhmmer run to extend the coordinates of each insertion by a user-specified number of nucleotides (250 by default) using `bedtools slop` from the BEDTools suite (Quinlan & Hall, 2010). After the extension, we extract a specified number of sequences (25 by default), selecting those with the highest bit-score from the `nhmmer` search, while ensuring no coordinate overlap between sequences.

The selected sequences are aligned using `MAFFT` (with parameters --`localpair --maxiterate 1000`)(Katoh & Standley, 2013), and the alignment is analyzed for the existence of multiple families with our family-splitting tool described in Submodule 2. If multiple families are detected, sequences corresponding to each family are split into individual files, and an HMMER model is generated with `hmmbuild` and then searched again with `nhmmer`, with the aim of compensating for the loss of sequences during the split. For each split family, we then extract the specified number of sequences from the new search (25 by default) and generate a new alignment with `MAFFT` using the same parameters.

After generating a new alignment for each family, or, if no other family was found, the module cleans up the alignment using the tool described in Submodule 1. In short, this tool generates multiple MSAs using various cleanup thresholds, which are used to construct HMM models. The models are searched in the genome using `nhmmer` to identify the best model, which is the one with the highest median bit-score.

To complete this module, we estimate whether each end of the model has grown in length. This is done by aligning the consensus of the new extended model to the consensus of the original model using MAFFT. Using Submodule 3, we analyze this alignment to determine which end of the alignment has grown in length and note the exact coordinates spanning the extended segment.

### Module 3: Side extension

The goal of this module is to extend each end of the model generated in Module 2. The input for this module consists of the alignment file generated by Module 2 and the coordinate file generated by Submodule 3. This module always starts by extending the 5’ end of the model first. This module generates an HMM model using `hmmbuild`, using the same parameters as described in Module 1. Then, the tool searches the entire model using `nhmmer` and selects only instances that overlap with the coordinate file generated by Submodule 3 in the previous module, thereby minimizing the risk of including fragments that might belong to other repeats. These coordinates are extracted and extended towards the 5’ end using `bedtools slop` using the same extension size used in Module 2.

From these extended coordinates, we extract the sequences and align them with `MAFFT`. After alignment, we determine whether multiple families are present using Submodule 2 and segregate them into separate alignment files. Then the module adds additional sequences to reach again the specified number (by defaults 25) as described in Module 2. After the MSA has been generated, the tool removes noisy regions of the alignment using the tool described in Submodule 1, and it selects the best cleaned MSA by generating an HMM with `hmmbuild` and searching the genome in the same manner as in Module 2.

The resulting MSA is then merged with the resulting MSA from Module 2 using `MAFFT’s --add` function (with parameters `--localpair --maxiterate 1000 --lexp -1.5 --lop 0.5`), and then gappy regions are removed by using the tool presented in Submodule 1. Coordinates of the extended region are determined by aligning the new consensus sequence with the previous one using MAFFT, and then passing the resulting MSA to Submodule 3.

After a round is finished, the pipeline determines whether there was a substantial increase (25% of the extension size by default). If there was a substantial increase, the next round of extension will use the output of Submodule 3 of this round as input. If there is no substantial increase, the tool shifts to extend the 3’ end of the consensus. As input, it uses the coordinates generated by Submodule 3 in Module 2, but the most recent 5’-end alignment serves as the template.

Once both ends have been extended, the tool takes the last successful round and passes the MSA and consensus sequence to the next module.

### Module 4: Model Polishing

This module takes the final alignment and resulting consensus sequence as inputs. Using the `hmmbuild` program, we generate an HMM model from the final alignment, which we then use with `nhmmer` to search the genome. We then use the output to generate a bed file containing the genomic insertion locations, the coordinates covered by the consensus, and the hit bit score. Then, we use the tools `bedmap` from `bedops` to concatenate the instances that are located in the same genomic neighborhood (which is 10 kbp by default), while retaining the information of the bit-score and coordinates of the repeat model covered by the insertion.

Using this bed file, the tool described in Submodule 4 appends sequences to a FASTA file until the desired coverage is reached (150 at every position by default) or the tool runs out of candidate sequences. Using this FASTA file, we generate a final MSA using MAFFT, which will then be cleaned with the Submodule 1 tool.

### Module 5: Reducing redundancy

Because some of the models might be duplicated as they were extended from two different sections of a repeat, we use CD-HIT-EST (Fu et al., 2012) to reduce redundancy of the repeat library, using the parameters proposed by (Goubert et al., 2022) (`-G 0 -b 500 -M 5000 -T 8 -c 0.8 -aS 0.8`).

### Submodule 1: MSA cleaner

We developed a tool to clean MSAs, recognizing that repeats can vary in the number of mutations. This tool takes an MSA in FASTA format as input and first removes positions with insufficient coverage (5 bases by default). After these regions have been removed, we proceed to calculate a score adapted from the Trident estimator, a score to quantify the quality of protein MSAs (Nguyen, 2011):

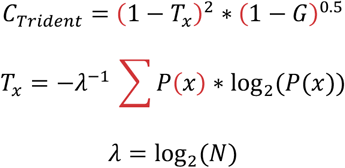

As in the original Trident estimator, Tx represents a normalized version of Shannon’s entropy, N is the number of sequences present in the alignment, P(x) is the probability of each nucleotide for a given position in the alignment, and G is the number of gaps present in each position. In essence, this estimator comprises two components: one that accounts for information content and the other that summarizes the number of gaps.

After calculating the information content for each position, we generate a density function and identify peaks in the distribution. The expectation is that columns corresponding to alignment noise will exhibit a distinct information distribution than homologous positions within the repeat. Using the peaks as thresholds, we remove positions whose median across windows (25, 15, 5, and 2 by default) is below the thresholds.

### Submodule 2: Family splitter

This submodule was designed to detect multiple families in an MSA. It starts by calculating the pairwise identity of each sequence present to generate a distance matrix. Across all distances, we generate a density function and identify peaks. In principle, if all sequences are alike to the same degree, the distribution will only contain one peak, even if the sequences are highly divergent. If two families are present within the same alignment, they will exhibit a bimodal distribution. If two or more peaks are present, the tool determines the distance between them; if this distance exceeds a set threshold (0.18 by default), the peaks are considered distinct. Finally, if there are at least two peaks, we calculate the midpoint between them and generate a phylogenetic tree, which is then cut at the height between the two peaks into subsets. The final output consists of several MSAs (in cases where there were multiple families).

### Submodule 3: Extract Coordinates

This submodule takes an MSA between the current and the last consensus sequence and outputs the coordinates at the 3’ and 5’ ends (in reference to the current consensus) where both sequences match for a specified number of positions (10 by default).

### Submodule 4: Window maker

This submodule aids the Polishing module by taking a bed file that contains the genomic coordinates of every single HMMER hit, as well as the bit score and the model’s coordinates. This tool starts by ordering the input file based on the bit-score and starts a loop where every entry’s genomic coordinate is extended by a set number of bases (10kbp by default) and checks for overlaps with other entries within this range. In cases of overlapping sequences, it merges the overlapping sequences with the original entry in order. If the sequence exceeds the length of the original sequence by a specified fraction (1.3 by default), the tool will output only the original entry. The tool will continue testing genomic windows until it reaches a specified per-base coverage (25 by default) or exhausts the sequence data.

### HEEA Implementation

The HEEA (HMMER-Extend-Extract-Align) algorithm is implemented inside the REE framework. It first searches each repeat model using HMMER, constructing the HMM model with `hmmbuild` using the parameters `--symfrac 0 --fragthresh 1.0 ––wnone`. Using the hmm model, we search for genomic instances with nhmmer, with default parameters (filtering out hits with an e-value < 1e-05). The tool orders the hits by bit score and samples non-overlapping hits for the extension (25 instances by default).

The instances’ coordinates are extended using `bedtools slop` (250 bp by default), and their sequences are extracted. The sequences are then aligned through `MAFFT` (with parameters `-- localpair --maxiterate 1000`). The resulting MSA is cleaned through the tools contained in Submodule 1. We then compare the resulting consensus with the original consensus or with the consensus generated in a previous round of extension, using the tool described in Submodule 3, to determine whether to proceed to another round of extension. If the consensus proceeds to another round, we reuse the genomic coordinates from the previous rounds and extend them by the same amount.

### Ancient Repeat Extension

Since ancient repeats are more fragmented and degenerated than younger repeats, we implemented an alternate pipeline to extend them using a recursive framework. To activate this mode within RRE, the user needs to use the `--AncientMode` flag.

The first difference between the conventional RRE approach and the ancient repeat extension is that the script described in Submodule 1 removes the peak with the lowest Trident estimator, rather than screening multiple peaks. This is because these repeats are often close to the signal-to-noise detection limit. The next difference is the removal of the family-splitting step in both the Central and Side extensions, as described in Submodule 2. Another significant change is that, instead of searching the entire repeat model, this approach searches only the previously extended sections, thereby maximizing the likelihood of capturing most sequences near the end of the model and removing the dependency on the combined MSA. Lastly, the final major change to this workflow is the use of the -- seed option in MAFFT instead of --add, as we observed that --add fails to incorporate the new round’s alignment at the right position in the combined MSA after a large number of rounds.

### RepeatMasker using HMMER models

Since RepeatMasker requires a GA (gathering) threshold to use HMMER as its search engine, the first goal is to calculate this threshold for each repeat model. We followed a similar approach to that established in Wheeler et al. (2012), by first calculating the bit score at which a false discovery rate (FDR) of 0.02% is achieved. To do this, we designed a Nextflow pipeline that takes a genome file in FASTA format and a file containing repeat-model alignments in Stockholm format (.stk). Next, the pipeline divides the genome into fixed-size segments (20 Mb by default). It generates a reverse but not complementary version of each segment, yielding forward (FWD) and reverse (REV) versions of the genome. At the same time, the tool generates HMM models from the .stk file and produces bundles of a specified number of models (20 by default).

After generating the genomic chunks and the HMM model bundles, we search all the models in both the FWD and REV versions of the genome with `nhmmer` with default parameters. After searching the hmm models across all genomic chunks, we determine the bit score at which the genomic hits from the REV genome constitute 0.02% of all hits, and we incorporate this GA threshold into a new HMM model.

To generate the final annotation, we search all models (which now include a GA threshold) with RepeatMasker using HMMER as the search engine and the FWD genome chunks as input. Since the FWD genome has been split into chunks, once all models have been searched we reconstitute the coordinates of the original genome in the RepeatMasker output file. Lastly, we use the corrected output file to mask the original genome and produce a RepeatMasker summary file.

